# *Neisseria gonorrhoeae* scavenges host sialic acid for Siglec-mediated, complement-independent suppression of neutrophil activation

**DOI:** 10.1101/2024.01.17.576097

**Authors:** Amaris J Cardenas, Keena S. Thomas, Mary W. Broden, Noel J. Ferraro, Constance M. John, Marcos M. Pires, Gary A. Jarvis, Alison K. Criss

## Abstract

Gonorrhea, caused by the bacterium *Neisseria gonorrhoeae* (Gc), is characterized by neutrophil influx to infection sites. Gc has developed mechanisms to resist killing by neutrophils that include modifications to its surface lipooligosaccharide (LOS). One such LOS modification is sialylation: Gc sialylates its terminal LOS sugars with cytidine-5’-monophosphate-*N*-acetylneuraminic acid (CMP-NANA) scavenged from the host using LOS sialyltransferase (Lst), since Gc cannot make its own sialic acid. Sialylation enables sensitive strains of Gc to resist complement-mediated killing in a serum-dependent manner. However, little is known about the contribution of sialylation to complement-independent, direct Gc-neutrophil interactions. In the absence of complement, we found sialylated Gc expressing opacity-associated (Opa) proteins decreased the oxidative burst and granule exocytosis from primary human neutrophils. In addition, sialylated Opa+ Gc survived better than vehicle treated or Δ*lst* Gc when challenged with neutrophils. However, Gc sialylation did not significantly affect Opa-dependent association with or internalization of Gc by neutrophils. Previous studies have implicated sialic acid-binding immunoglobulin-type lectins (Siglecs) in modulating neutrophil interactions with sialylated Gc. Blocking neutrophil Siglecs with antibodies that bind to their extracellular domains eliminated the ability of sialylated Opa+ Gc to suppress oxidative burst and resist neutrophil killing. These findings highlight a new role for sialylation in Gc evasion of human innate immunity, with implications for the development of vaccines and therapeutics for gonorrhea.

**IMPORTANCE:** *Neisseria gonorrhoeae,* the bacterium that causes gonorrhea, is an urgent global health concern due to increasing infection rates, widespread antibiotic resistance, and its ability to thwart protective immune responses. The mechanisms by which Gc subvert protective immune responses remain poorly characterized. One way *N. gonorrhoeae* evades human immunity is by adding sialic acid that is scavenged from the host onto its lipooligosaccharide, using the sialyltransferase Lst. Here, we found that sialylation enhances *N. gonorrhoeae* survival from neutrophil assault and inhibits neutrophil activation, independently of the complement system. Our results implicate bacterial binding of sialic acid-binding lectins (Siglecs) on the neutrophil surface, which dampen neutrophil antimicrobial responses. This work identifies a new role for sialylation in protecting *N. gonorrhoeae* from cellular innate immunity, which can be targeted to enhance the human immune response in gonorrhea.

## INTRODUCTION

*Neisseria gonorrhoeae* (Gc) causes the bacterial sexually transmitted infection gonorrhea. Gc is designated as an urgent threat level pathogen by the Centers for Disease Control and Prevention, with approximately 82.4 million cases each year globally (1). Factors contributing to the prevalence of gonorrhea include increasing resistance to antibiotics, resistance to soluble and cellular innate immune components, and the lack of protective immunity elicited by prior infection (2, 3). Finding new approaches to treat or prevent Gc infection is of utmost importance.

The most abundant surface component on Gc is lipooligosaccharide (LOS). The length and composition of LOS is regulated by *lgt* (LOS glycosyltransferase) genes. Phase variation of *lgt* genes results in varying oligosaccharide (OS) compositions within a Gc population, including during infection (4–7). Gc isolated from uncomplicated urethral infection predominantly produce LOS that can be sialylated. Sialylation is the incorporation of sialic acid onto the alpha and/or beta chain terminal galactose of LOS, catalyzed by Gc LOS sialyltransferase (Lst) (8–10). Gc cannot synthesize sialic acid and instead scavenges CMP-NANA from the host (11). Lst is constitutively expressed (12, 13), is required for optimal genital tract infection (14–16), and uses diverse forms of CMP-sialic acid (17).

LOS sialylation is a form of molecular mimicry by which Gc is thought to evade immune recognition (18, 19). Sialylation was originally characterized as conferring “unstable” serum resistance on Gc recovered from urethral secretions, but not after lab passage (11, 20, 21). Sialylation has since been shown to inhibit all three complement activation pathways by decreasing C1 engagement and C4b deposition, and increasing recruitment of factor H on Gc (17, 22, 23). Sialylated glycans enable the immune system to discriminate self from non-self, achieved in part through sialic acid-binding immunoglobulin-type lectins (Siglecs) (24–27). Siglecs are produced by most immune cells, including neutrophils, which are the predominant immune cells recruited in Gc infections and constitute the purulence in gonorrheal secretions (28). Human neutrophils express Siglec-5, -9, and -14. Siglec-5 and -9 contain cytoplasmic immunoreceptor tyrosine-based inhibitory motifs (ITIMs) that recruit SH2 domain-containing phosphatases to inhibit cellular activation (29–34). In contrast, Siglec-14 associates with the adapter protein DAP12, which contains an immunoreceptor tyrosine-based activating motif (ITAM) that once phosphorylated, recruits Syk kinase to activate downstream signaling (35, 36). The genes that encode Siglec-5 and Siglec-14 are adjacent and considered paired receptors, since the proteins share almost complete sequence identity in their binding domains and similar glycan binding preferences (35, 37). 10-70% of individuals in certain racial/ethnic groups harbor a *SIGLEC14/5* fusion gene where Siglec-14 is not expressed, but Siglec-5 is (38). Thus, on balance, signaling through neutrophil Siglecs dampens inflammatory signaling. Sialylated Gc has been reported to interact with recombinant extracellular domains of these Siglecs (39).

Gc expresses numerous gene products that defend against neutrophil antimicrobial activities

(28). Gc also varies its ability to interact non-opsonically with neutrophils through phase-variable expression of opacity-associated (Opa) proteins (40). Most Opa proteins bind one or more carcinoembryonic antigen-related cell adhesion molecules (CEACAMs). In particular, binding to the granulocyte-restricted CEACAM3 elicits phagocytosis, reactive oxygen species production, and granule release (41–47).

In this study, we unexpectedly found that sialylation promotes Gc survival from human neutrophils in a complement-independent manner. Our results implicate Siglecs in dampening the neutrophil activation that is elicited by CEACAM-binding Opa proteins, bringing new insight into the ways that Gc modulates soluble and cellular human innate immunity to persist in its obligate human host.

## RESULTS

### Gc Lst incorporates sialic acid onto LOS, which is retained upon infection of neutrophils

To investigate the role of LOS sialylation on direct, nonopsonic Gc interactions with neutrophils, we first assessed Lst- and CMP-NANA-dependent sialylation of Opa-expressing (Opa+) Gc. The wild-type (WT) Lst parent for these studies was a derivative of strain FA1090 1-81-S2 that constitutively expresses the OpaD protein in an otherwise Opa-negative background, hereafter termed OpaD (48). OpaD binds CEACAMs 1 and 3 on human neutrophils, leading to rapid phagocytic killing (48, 49). This strain produces the sialylatable lacto-*N*-neotetraose (LNnT) alpha chain and lactose beta chain (50). OpaD was transformed with insertionally inactivated *lst* to generate an isogenic mutant that cannot be sialylated (OpaDΔ*lst*) (15). Bacteria were incubated with CMP-NANA or vehicle control, then washed to remove residual sialic acid and processed for downstream analysis.

Monoclonal antibody (mAb) 6B4 binds the LNnT epitope of the alpha chain of *Neisseria* LOS; α2,3 sialylation of its terminal galactose prevents 6B4 binding (8, 51). By Western blot, 6B4 reacted with lysates of OpaD, and binding was lost when OpaD was incubated with CMP-NANA (**Fig 1A**). 6B4 reacted with OpaDΔ*lst* lysates with or without CMP-NANA (**Fig 1A**). To analyze sialylation on individual bacteria, we used imaging flow cytometry with 6B4. The fluorescence index of OpaD incubated with CMP-NANA was significantly lower than OpaD without CMP-NANA and OpaDΔ*lst* with or without CMP-NANA; these three were statistically indistinguishable (**Fig 1B-C**).

**Figure 1.**
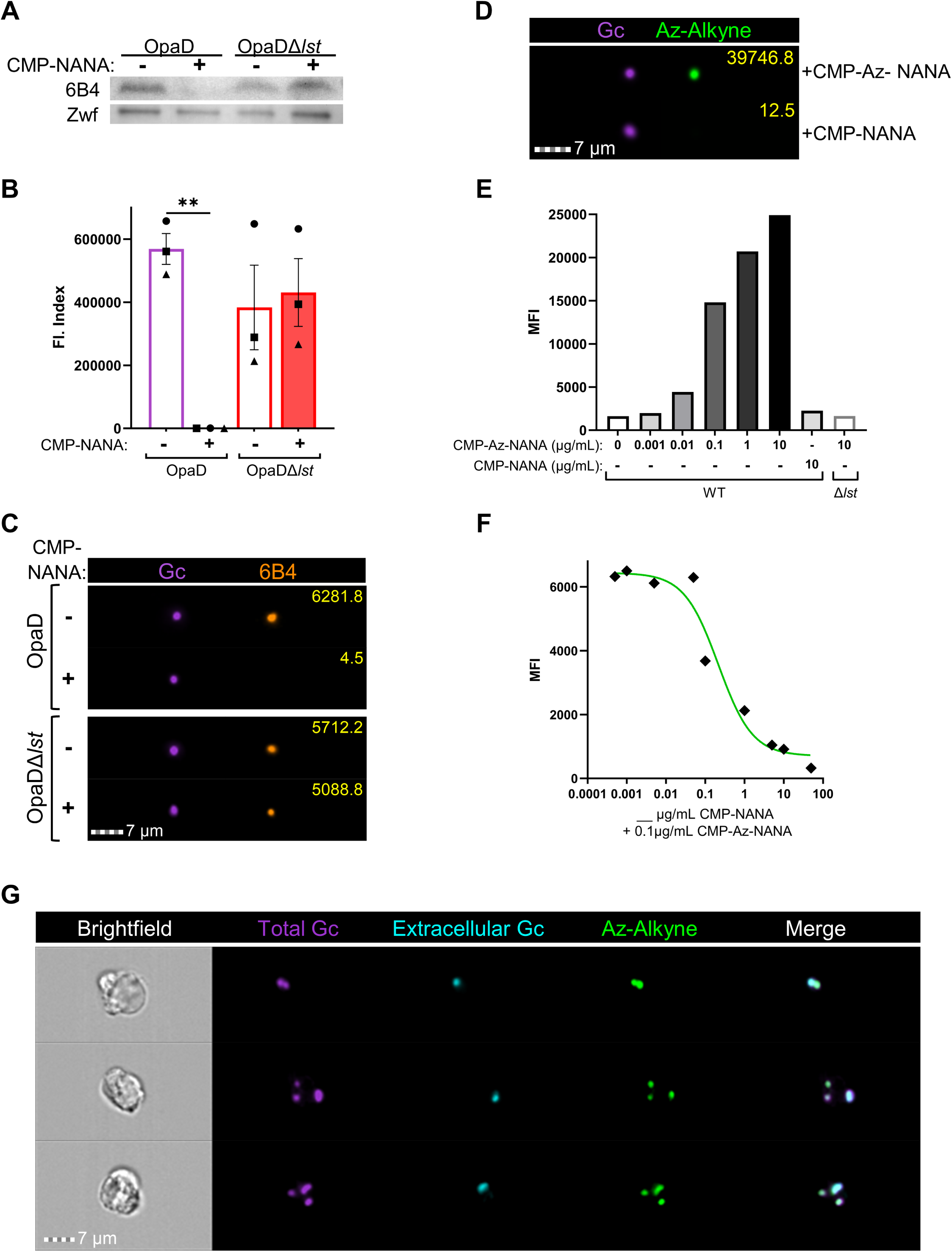
Analysis and quantification of Lst-dependent sialylation of gonococcal lipooligosaccharide. (**A**) OpaD and OpaDΔ*lst* were grown with 50µg/mL CMP-Neu5Ac (NANA) (+) or vehicle (Veh) (-). Bacterial lysates were Western blotted with mAb 6B4, or anti-Zwf as loading control, followed by HRP-conjugated secondary antibodies. Blot is representative of n=3 biological replicates. (**B-C**) OpaD and OpaDΔ*lst* were incubated with CMP-NANA or vehicle as in **A**, then stained with Tag-IT Violet (TIV). Bacteria were incubated with 6B4 followed by AlexaFluor647-coupled secondary antibody, fixed, and analyzed by imaging flow cytometry. (**B**) Average fluorescence index (Fl: mean fluorescence intensity (MFI) x percent positive) from n=3 biological replicates with 20,000 events collected per condition; each symbol indicates one matched biological replicate. Statistical analysis by one-way ANOVA with Tukey’s multiple comparisons test. **=p≤0.01. (**C**) Representative images for the indicated condition; AF647-6B4 signal is false-colored orange, and yellow numbers indicate the object’s fluorescence intensity value. (**D-G**) Gc were incubated with either CMP-NANA and/or CMP-Az-NANA at 100ng/mL. Bacteria were stained with TIV, fixed, then subjected to copper catalyzed click chemistry azide-alkyne cycloaddition using fluorescein isothiocyanate (FITC)-alkyne. (**D**) Representative images of CMP-NANA and CMP-Az-NANA sialylated OpaD, each of which was subjected to alkyne-FITC cycloaddition. Yellow numbers are as in **C**. (**E**) OpaD or OpaD*Δlst* were incubated with indicated concentrations of CMP-Az-NANA or CMP-NANA. The MFI of each condition was quantified from flow cytometry, n=1. (**F**) WT OpaD was incubated with 0.1µg/mL of CMP-Az-NANA along with the indicated concentration of unmodified CMP-NANA. Results are graphed as the MFI of each condition and fit with a non-linear regression line. (**G**) OpaD sialylated with CMP-Az-NANA was stained with TIV and used to infect IL-8 treated, adherent primary human neutrophils for 30 min. After fixation, extracellular Gc were labeled with anti-PorB and AF647-coupled secondary antibody (false-colored cyan) before permeabilization with 0.1% saponin followed by copper click chemistry. Representative images of infected neutrophils with intracellular (TIV+AF647-) and extracellular (TIV+AF647+) Gc with azide-sialylated LOS that is clicked with the alkyne-FITC probe (FITC+).

Two other sialylation sites on LOS have been reported in Gc: the P^k^-like alpha chain (18, 52) and the lactose-bearing beta chain (50, 53), both through α2,6 linkage. To directly detect Lst-dependent sialylation on Gc LOS, independent of 6B4 exclusion, we applied copper click chemistry using azido-labeled sialic acid, followed by alkyne-FITC cycloaddition (54–57). OpaD incorporated CMP-Azido (Az)-NANA in a Lst-dependent manner (**Fig 1D-E**), which was displaced in competition with unmodified CMP-NANA at approximately equimolar concentration (**Fig 1F**). Using Gc preincubated with CMP-Az-NANA, we monitored the retention of sialylation on OpaD in the presence of neutrophils. Representative images collected at 30 min post-infection showed alkyne-FITC labeled bacteria that were both intracellularly and extracellularly attached to neutrophils (**Fig 1G**). Thus, imaging flow cytometry, in combination with immunofluorescence and click chemistry, can be used to track and quantify Gc LOS sialylation alone and in cells.

### Lst-dependent sialylation of Opa+ Gc dampens human neutrophil activation and protects Gc from killing by neutrophils

We explored the contribution of Gc Lst-dependent sialylation to neutrophil activation. First, the release of reactive oxygen species (ROS) was measured. As previously reported, nonsialylated OpaD elicited robust neutrophil ROS production (**Fig 2A-B**) (42, 45, 48). Sialylation significantly reduced the magnitude of the OpaD-elicited oxidative burst; ROS production was inversely correlated with the concentration of CMP-NANA for sialylation (**Fig 2A-B**). Sialylation similarly reduced the oxidative burst in response to other Opa+ bacteria (**Fig 2C, Fig S1A**). OpaDΔ*lst* elicited neutrophil ROS production regardless of CMP-NANA addition, and was indistinguishable from nonsialylated OpaD (**Fig 2D, Fig S1B**).

**Figure 2.**
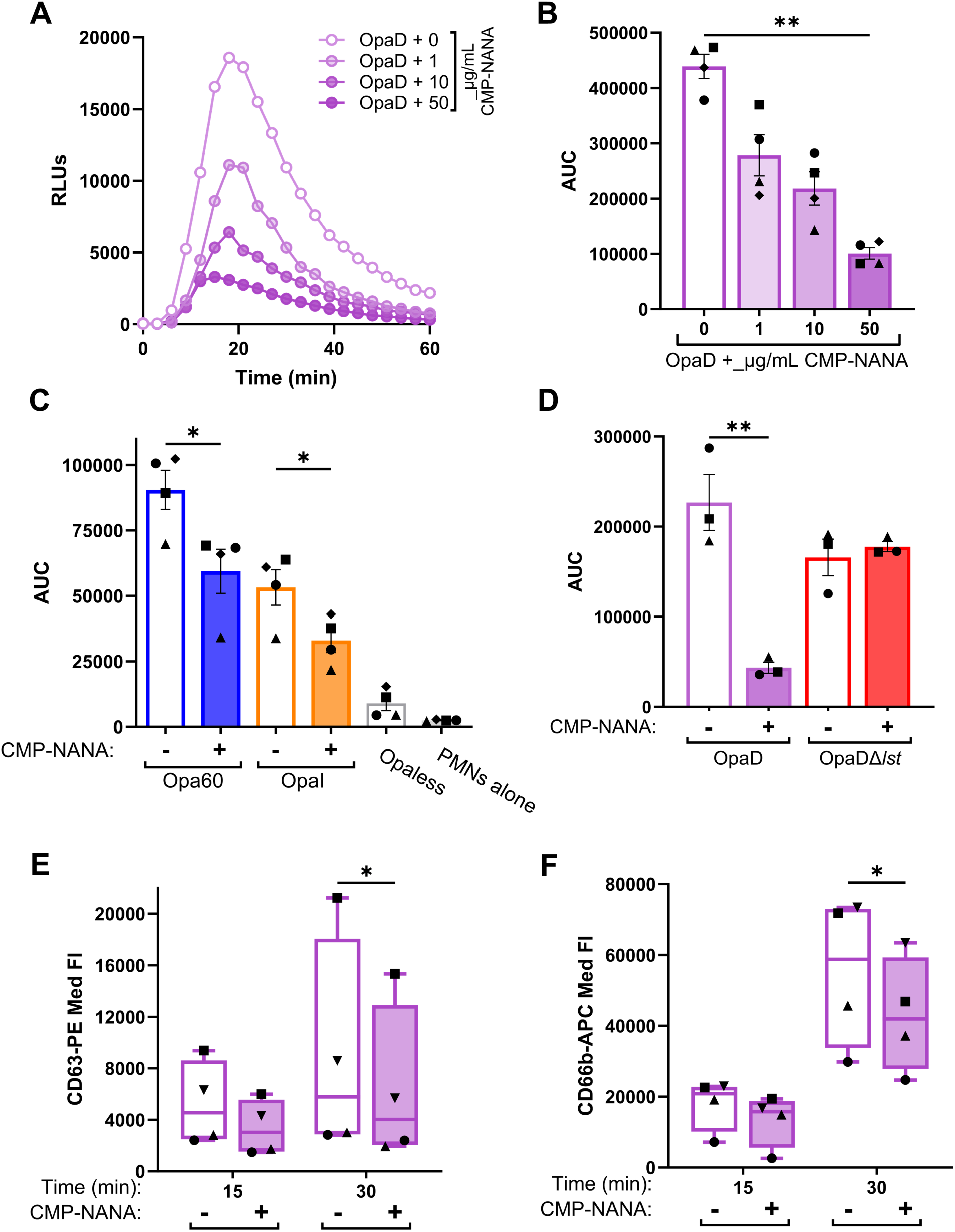
Sialylation dampens neutrophil activation in response to Opa+ bacteria. (**A-B**) OpaD incubated with the indicated concentrations of CMP-NANA or vehicle (0 µg/mL) were exposed to primary human neutrophils in the presence of luminol at an MOI of 100. Neutrophil oxidative burst was measured as relative light units (RLUs) of luminol-dependent chemiluminescence every 3 min over 1 h. (**A**) displays one representative graph of n=4; (**B**) is the average ± SEM area under the curve (AUC) for each condition across replicates (symbols indicate matched biological replicates). (**C**) Opa60 Gc (blue) or OpaI Gc (orange) were incubated with 50 µg/mL CMP-NANA (+) or vehicle (-) before neutrophil exposure; AUCs of n=4 biological replicates. Non-stimulatory Opaless Gc (dark grey) exposed neutrophils or neutrophils (PMNs) in luminol alone (light grey) serve as negative controls. (**D**) WT (purple) or Δ*lst* (red) OpaD were treated with or without CMP-NANA then added to neutrophils as above; AUCs of n=3 biological replicates. Statistical analyses by one-way ANOVA with Tukey’s multiple comparisons test. *=p≤0.05; **=p≤0.01. (**E-F**) OpaD treated with CMP-NANA or vehicle as above were added to adherent, IL-8 primed neutrophils for 15 or 30 min at an MOI of 1. Neutrophils were stained for viability (Zombie Near Infrared) and with a PE-coupled antibody against primary granule protein CD63 (**E**) and an APC-coupled antibody against secondary granule protein CD66b (**F**) on the cell surface. After fixation, cells were analyzed via spectral flow cytometry. Data are presented as median fluorescence (Med Fl) of PE+ (**E**) and APC+ (**F**) live cells, gated using unstained and isotype controls. Results are from n=4 biological replicates (symbol matched). Statistical analyses were performed by two-way ANOVA with Šídák’s multiple comparisons test. *=p≤0.05.

ROS does not directly contribute to neutrophil antigonococcal activity (58–60), ROS production requires intracellular signaling events and granule release (61). Neutrophil granules contain antimicrobial components, some of which are bactericidal for Gc (62–64). Thus, we analyzed the effect of sialylation on the ability of OpaD to stimulate exocytosis of primary and secondary neutrophil granules as measured by CD63 and CD66b surface presentation, respectively (42). Sialylation of Gc significantly reduced neutrophil exocytosis of primary (**Fig. 2E**) and secondary (**Fig 2F**) granules at 30 min post-infection. Together, these data indicate sialylation reduces the ability of Gc to activate neutrophils.

We next tested the contribution of Lst-dependent sialylation to bacterial survival in the presence of adherent, IL-8 primed primary human neutrophils without addition of human serum (complement). Sialylated OpaD had a statistically significant increase in survival over nonsialylated bacteria at 15 and 30 min postinfection (**Fig 3A**). OpaDΔ*lst* phenocopied nonsialylated OpaD at these times, showing that both sialic acid and Lst were required for the increased survival of OpaD exposed to neutrophils. By 60 and 120 min post-infection, survival of sialylated OpaD was not statistically different from nonsialylated Gc (**Fig 3A**). Sialylation also significantly increased the survival from neutrophils of CEACAM-binding Opa60+ Gc, through 2 h infection (**Fig 3B**). However, survival of Opaless Gc from neutrophils was unaffected by sialylation, which fits with our previous reports that Opa-negative Gc persist in the presence of neutrophils (48, 59, 65) (**Fig 3C**). These results highlight a role for Lst-dependent sialylation in survival of Opa+ Gc from neutrophils.

**Figure 3.**
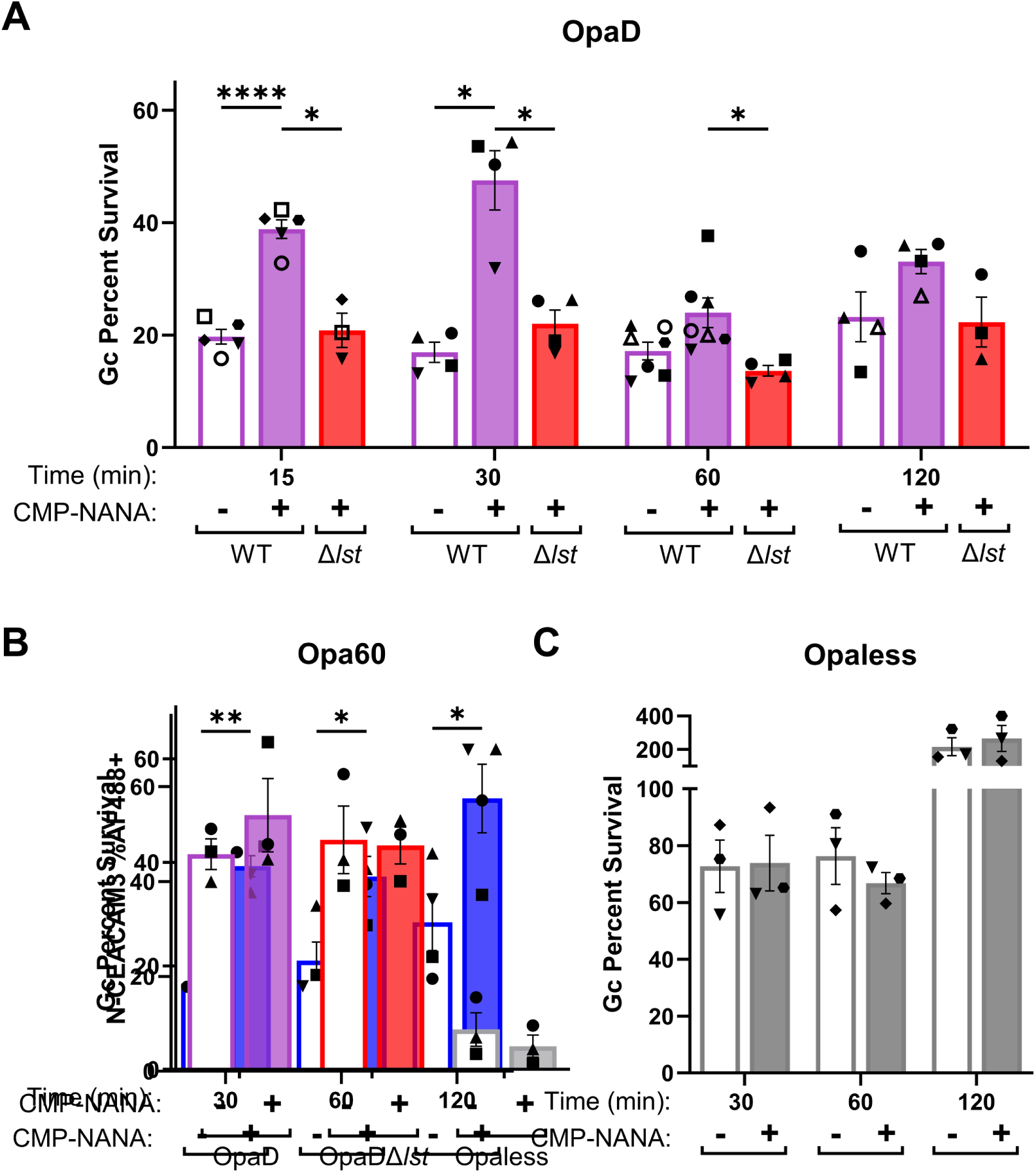
Sialylated, Opa+ bacteria have a survival advantage at early times of neutrophil challenge. Adherent, IL-8 primed neutrophils were infected with sialylated (filled) or nonsialylated (empty) (**A**) OpaD (purple) or OpaDΔ*lst* (red), at a MOI of 1. Gc viability was determined by the enumeration of CFUs from lysed neutrophils at the indicated times post-infection, expressed as a percentage of CFU at 0 min. (**B-C**) Sialylated and nonsialylated Opa60 Gc (blue) (**B**) and Opaless Gc (grey) (**C**) were exposed to neutrophils and bacterial viability measured as in **A**. Results are presented as the average ± SEM for n≥3 (**A**) n=4 (**B**) or n=3 (**C**) biological replicates, matched by symbol within each data set. Statistical comparisons were by mixed effect analysis (**A**) or two-way ANOVA (**B-C**) with Holm-Šídák’s multiple comparisons test with the following pairwise significances: *=p≤0.05; **=p≤ 0.01; ****=p≤0.0001.

### Sialylation of LOS does not directly affect OpaD-CEACAM3 interactions

Given the importance of CEACAM3 to phagocytic killing of Opa+ Gc by neutrophils, we tested the hypothesis that Gc sialylation disrupted OpaD-CEACAM3 engagement. First, sialylated or nonsialylated OpaD was incubated with a Glutathione S-transferase (GST)-tagged N-terminus of human CEACAM3 (N-CEACAM3), and CEACAM3 binding was measured by flow cytometry using anti-GST fluorescence (66). Lst-dependent sialylation did not significantly affect OpaD binding of N-CEACAM3 (**Fig 4A**). Sialylation also did not affect interaction of OpaD with N-CEACAM1 (**Fig S2**). Opaless Gc, which does not bind CEACAMs, served as negative control. Second, association of sialylated or nonsialylated OpaD with human CEACAM-3 transfected CHO cells was measured by imaging flow cytometry. Lst-dependent sialylation did not significantly affect bacterial association with hCEACAM3-CHO cells (**Fig 4B**). These results indicate that contrary to our hypothesis, sialylation does not affect Opa-CEACAM engagement.

**Figure 4.**
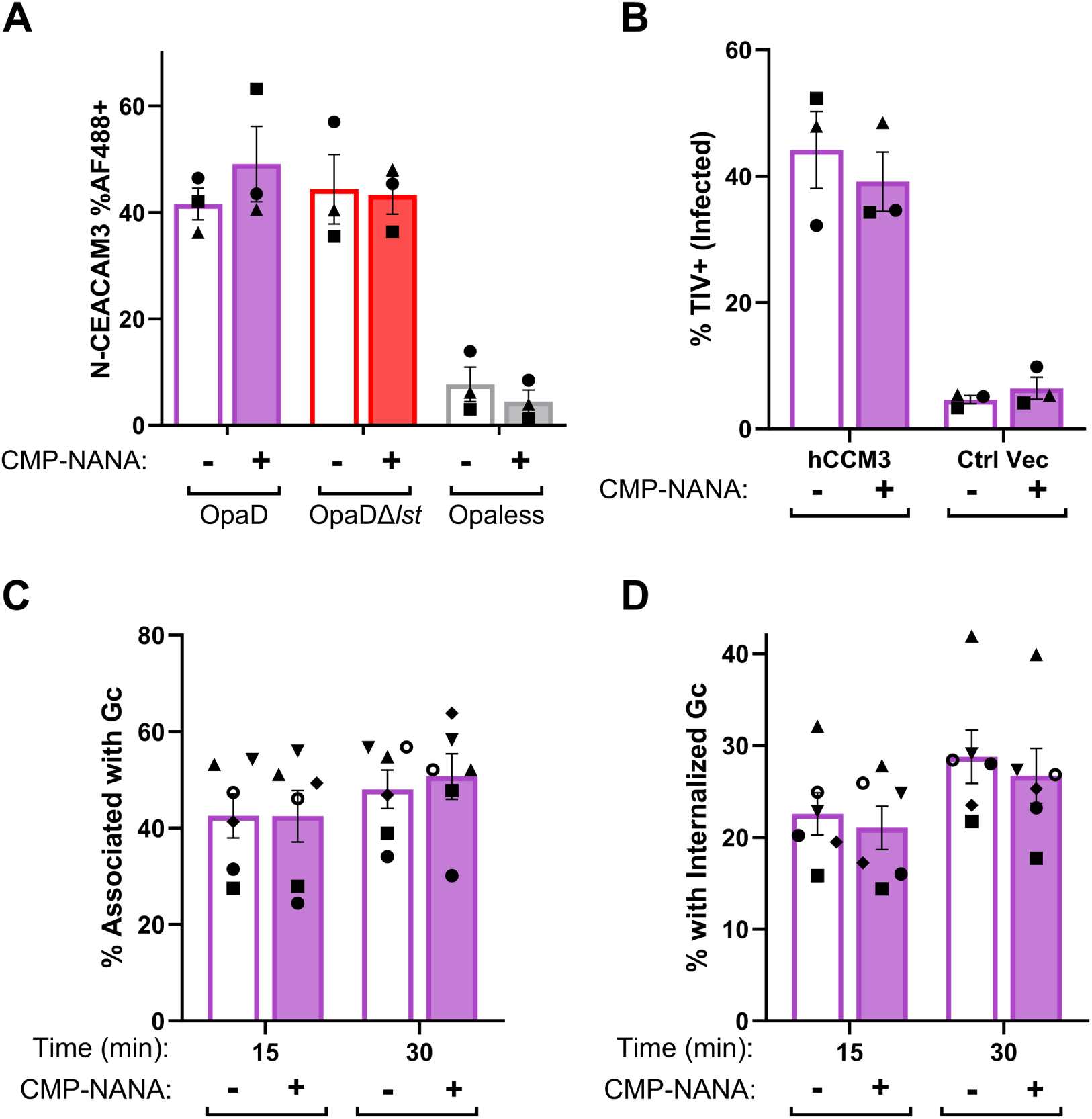
Sialylation does not affect binding of OpaD bacteria to CEACAM-3. (**A**) OpaD grown with CMP-NANA or vehicle were incubated with GST-tagged recombinant N-terminal domain of CEACAM-3 (NCEACAM-3). Binding of NCEACAM-3 to Gc was detected using a mouse anti-GST antibody, followed by a goat anti-mouse IgG AF488-conjugated antibody. Gc were fixed, stained with DAPI, and analyzed by imaging flow cytometry, to calculate the percent of singlet Gc that are AF488+. Opaless Gc that does not bind CEACAM is shown as a negative control (grey). (**B**) CHO cells transfected with human CEACAM3 (hCCM3) or empty control vector (Ctrl Vec) were infected with TIV-labeled OpaD, treated with CMP-NANA or vehicle. After 30 min, cells were collected, stained with a pan-CEACAM antibody followed by AF555-conjugated secondary antibody, and fixed. The percent of singlet CHO cells that are TIV+ (infected) was calculated using imaging flow cytometry. Results are from n=3 biological replicates (symbol matched). Statistical comparisons were by two-way ANOVA with Tukey’s multiple comparisons test; not significant. (**C-D**) OpaD with or without sialylated LOS were labeled with TIV and used to infect adherent, IL-8 primed neutrophils. At the indicated time points, infected cells were fixed and extracellular Gc were detected with an anti-PorB antibody, followed by an AF488-coupled secondary antibody, without permeabilization. Cells were then analyzed using imaging flow cytometry to report the percent of neutrophils with associated (%TIV+) Gc (**C**) or internalized (%TIV+AF488-) Gc (**D**). Results are from n=4 biological replicates (symbol matched). Statistical comparisons were by two-way ANOVA with Tukey’s multiple comparisons test; not significant.

In a previous report, sialylation decreased the adherence of Opa+ Gc to neutrophils in suspension, but this effect was lost over time (67). We thus hypothesized that the mechanism by which sialylated Gc resisted killing by neutrophils was due to decreased association with and/or internalization by these immune cells. To test this hypothesis, we used imaging flow cytometry (68), in which extracellular Gc were discriminated from intracellular bacteria by accessibility of a mAb against FA1090 PorB1b. Binding of this antibody to Gc was unaffected by sialylation (**Fig S3**). Sialylation did not significantly affect OpaD association with (**Fig 4C**) or internalization by (**Fig 4D**) neutrophils through 30 min post-infection, when effects were measured by ROS production and bacterial survival. We conclude that sialylation does not directly affect Opa+ Gc interaction with neutrophil CEACAMs.

### Sialylated Gc bind immunoregulatory Siglecs to dampen neutrophil activation and antigonococcal activity

The inhibitory effect of Gc sialylation on neutrophil activation and not phagocytosis, along with work from other groups (37, 39, 69, 70), led us to investigate the contribution of sialic acid immunoglobulin-type lectins (Siglecs). Using imaging flow cytometry, we confirmed the presence of Siglec-9 and Siglec-5/14 on the surface of primary human neutrophils (**Fig 5A**) compared to unstained (**Fig S4A**). Staining ranged from small puncta to rings around the cell periphery, though there was minimal overlap in Siglec-9 vs. Siglec-5/14 localization (**Fig 5B**).

**Figure 5.**
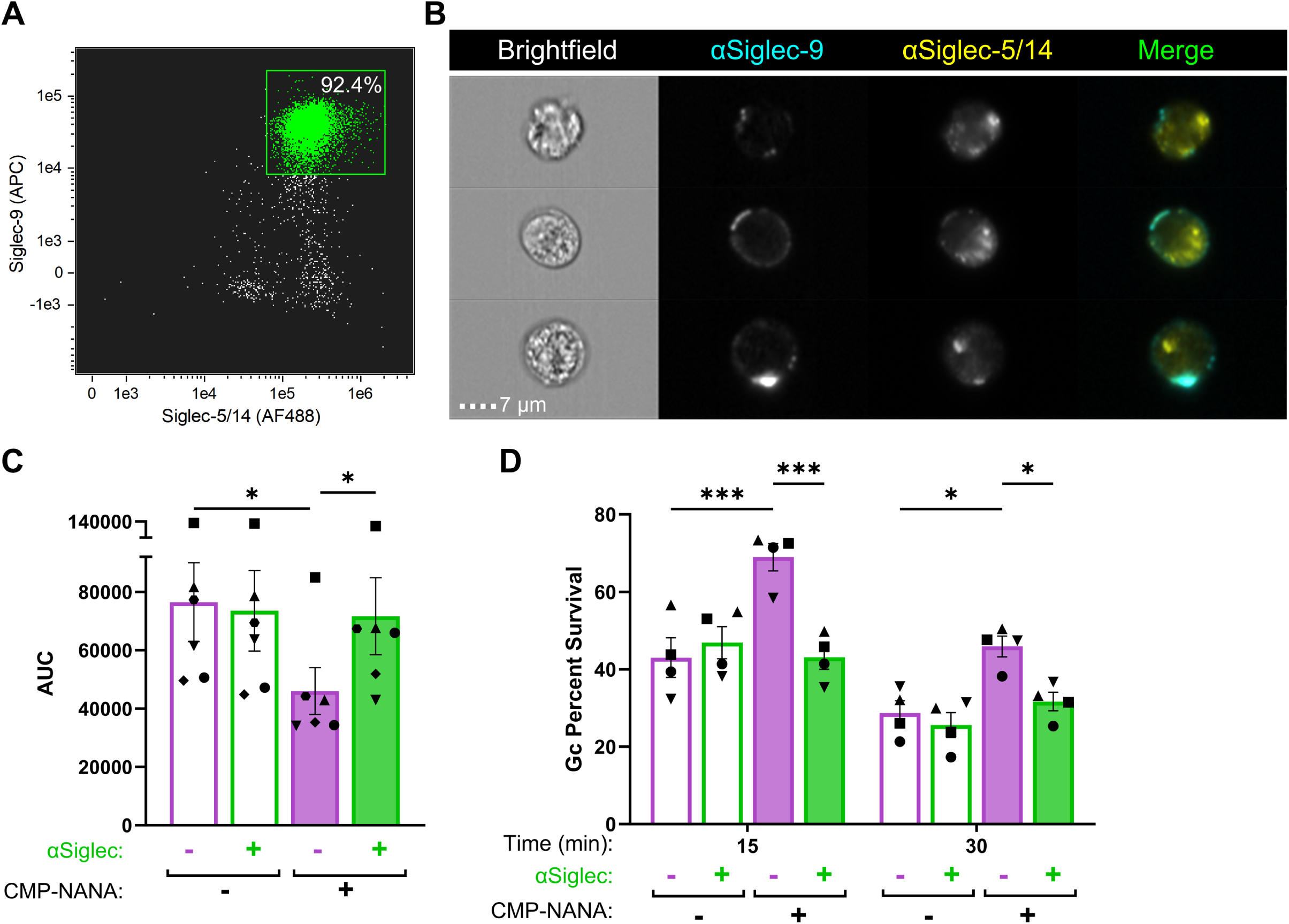
Blockade of neutrophil Siglec-9 and Siglec-5/14 reverses the ability of sialylated gonococci to suppress neutrophil activation and antibacterial activity. (**A-B**) Uninfected and adherent, IL-8 primed human neutrophils were incubated with antibodies that recognize human Siglec-9-APC and/or Siglec-5/Siglec-14-AF488 antibodies, without cell permeabilization before fixation and processing via imaging flow cytometry. (**A**) Dot plot of focused singlet neutrophils with anti-Siglec-9(APC) and anti-Siglec-5/14(AF488). X axis indicates AF488 fluorescence intensity and y axis indicates APC fluorescence intensity. Double positive (green) gate based on single stains; number within gate is the percent of the stained condition. (**B**) Representative images of neutrophils stained with anti-Siglec-9-APC (cyan) and anti-Siglec-5/14-AF488 (yellow); colored fluorescence in merged. Results are representative of n=3 biological replicates by spectral flow cytometry and confocal microscopy (not shown) and demonstrate n=1 replicate with imaging flow cytometry. (**C-D**) Primary human neutrophils were incubated with anti-human Siglec-9 and Siglec-5/-14 antibodies (green) or media alone (purple) for 30 min. (**C**) Neutrophils were then exposed to sialylated (filled) or nonsialylated (empty) OpaD at an MOI of 100 in the presence of luminol. ROS production was measured over 1 h by luminol-dependent chemiluminescence as RLUs. Graphed data of AUCs for each condition across n=6 biological replicates (symbol matched). Statistical comparisons by one-way ANOVA with Tukey’s multiple comparisons test. *=p≤0.05. (**D**) Neutrophils were treated with anti-human Siglec-9 and Siglec-5/-14 antibodies (green) or media alone (purple) as in **C**. Cells were then infected with sialylated (filled) or nonsialylated (empty) Gc at an MOI of 1. Survival of Gc as a percent of the bacteria at time 0 min, as measured as in Fig 3. Statistical analyses by two-way ANOVA with Holm-Šídák’s multiple comparisons test. n=4 biological replicates (symbol matched); *=p ≤ 0.05, ***p<0.0001.

We hypothesized that sialic acid-Siglec engagement interfered with Opa-CEACAM binding to dampen the neutrophil response to Opa+ Gc. To test this, neutrophils were incubated with blocking antibodies against the extracellular domains of Siglec-9 and -5/14 (71), then exposed to sialylated or nonsialylated OpaD, and ROS production was measured. As seen in **Fig 2**, sialylation significantly reduced the neutrophil oxidative burst in response to OpaD when isotype control antibodies were added (purple bars, **Fig 5C**). Addition of anti-Siglec antibodies significantly enhanced the burst elicited by sialylated OpaD (solid green vs. solid purple, **Figs 5C** and **S4B**), restoring it to levels that were no different from unsialylated OpaD. The restoration of ROS in the presence of Siglec-blocking antibodies was specific to sialylated Gc, as nonsialylated OpaD induced similar levels of ROS whether or not blocking antibodies were present (green outline vs. purple outline, **Fig. 5D**). These data implicate Siglecs in the ability of sialylated Opa+ Gc to reduce neutrophil activation.

The dampened ROS release in response to sialylated Opa+ Gc led us to examine if blocking Siglecs restored the ability of neutrophils to control Opa+ bacteria. Neutrophils were incubated with blocking antibodies against Siglec-9 and -5/14 or left untreated, then exposed to sialylated or nonsialylated OpaD. Addition of anti-Siglec antibodies significantly reduced the survival of sialylated OpaD after infection of neutrophils at 15 and 30 min, to a level equivalent to nonsialylated OpaD (**Fig 5D**). Nonsialylated OpaD were recovered similarly from neutrophils whether or not anti-Siglec antibodies were present, indicating that the effect of Siglecs on Gc-neutrophil interaction is specific to sialylated Gc.

From these results, we conclude that Gc uses sialylation of its LOS in a Siglec-mediated, complement-independent manner to impede neutrophil activation and antigonococcal responses, thereby promoting bacterial survival during infection.

## DISCUSSION

Gc sialylation of LOS by Lst is crucial for complement resistance and pathogenesis *in vivo* (10). This study reveals an unexpected, complement-independent role of sialylation: its ability to restrain neutrophil activation in response to Opa+ Gc, improving Gc survival upon neutrophil challenge. Sialylation reduced granule mobilization and release of both oxidative and nonoxidative species in response to Opa+ bacteria. Unexpectedly, sialylation of the LOS did not markedly affect Opa-CEACAM engagement. Instead, by blocking the immunoregulatory Siglecs on neutrophils, sialylated Opa+ Gc could no longer suppress neutrophil activation or killing capacity. These data suggest the exploitation of neutrophil self-associated molecular pattern recognition by Gc through LOS sialylation by Lst. Our findings add to the increasing understanding of how pathogens co-opt host factors to impair immune cell activation (72).

This study focused on Gc interactions with neutrophils, the main cell type found in human infectious secretions. We used human neutrophils that were adherent and treated with IL-8, mimicking post-migration behavior in these terminally differentiated, short-lived primary cells (59). Work by our group and others has investigated how Gc expression of phase-variable Opa proteins affects its survival from neutrophils (40–42, 45, 48, 63, 73). Most Opa proteins bind selected human CEACAMs. In particular, neutrophils and other granulocytes express CEACAM3, which contains an ITAM that can recruit Src family and Syk tyrosine kinases to drive signaling pathways leading to phagocytosis, granule release, and ROS production (40). Given the negative charge conferred by sialic acids, we hypothesized that LOS sialylation would impair the phagocytic activity of neutrophils towards Gc. However, we found no significant difference in the ability of sialylated Opa+ Gc to bind the soluble N-domain of CEACAM, CEACAM-expressing cells, or primary human neutrophils. While our study only focused on Opa+ Gc that bind CEACAMs 1 and 3, future work can explore how sialylation affects other modes of interaction between Gc and neutrophils, including antibody- and complement-mediated association.

Siglecs help the innate immune system to distinguish between non-self and self-associated molecular patterns (26). Many immune cell Siglecs contain cytoplasmic ITIM domains that transduce inhibitory signals to dampen inflammation (24). Other sialylated bacteria, including *N. meningitidis* and Group B *Streptococcus*, have been shown to manipulate cell activation via Siglec engagement (27, 69), and Gc can bind human Siglec-Fc chimera proteins (39). Consistent with published findings, we detected Siglec-9 and Siglec-5/14 on the human neutrophil surface (29, 32, 37). Moreover, blocking Siglecs reversed the inhibitory activity of sialylation on neutrophil responses to Gc, uncovering a new role for Siglecs in thwarting neutrophil antigonococcal responses. Interestingly, human Siglec-Fc chimeras have also been reported to bind PorB on the surface of unsialylated Gc, which was enhanced by production of a shorter, nonsialylatable LOS (39). Whether nonsialylated Gc interact with Siglecs on neutrophils and the consequences of such interaction remain to be explored. Unlike Siglec-5 and Siglec-9, Siglec-14 does not have a cytoplasmic domain; instead, it associates with the ITAM-bearing DAP12 (35, 37). Siglec-5 and Siglec-14 have almost identical ligand binding domains due to ongoing gene conversion and are considered paired receptors, possibly to balance immune responses to pathogens (24, 35, 38). In a study of genetic variations in Siglecs among a human cohort with a high burden of gonorrhea, uninfected individuals were more likely to produce Siglec-14, though not reaching statistical significance (39). Future work can examine if neutrophils from individuals with or without Siglec-14 respond differently to sialylated Gc.

The localization of Lst has recently been updated from an outer membrane protein to the inner face of the cytoplasmic membrane, where the OS is assembled (74, 75). Unlike the closely related microbe *N. meningitidis*, Gc cannot endogenously synthesize CMP-NANA and must scavenge it from the extracellular environment for sialylation. Lst has both α2,3 (alpha chain LNnT) and α2,6 sialyltransferase activity (alpha chain P^k^-like OS and beta chain lactose) and has been shown to use different CMP-sialic acids as substrates (8, 17, 52, 53). This promiscuity suggests that sialic acid availability rather than enzymatic activity drives the dynamics of LOS sialylation. We exploited this feature of Lst to add azido-labeled CMP-NANA in the form of N-acetylneuraminic acid (Neu5Ac) to directly track sialylation on the Gc surface. Other Lst substrates include CMP-Neu5Gc, which is missing in humans due to the evolutionary loss of the responsible hydrolase (76), and CMP-legionaminic acid and CMP-ketodeoxynonulosonic acid, which do not confer complement-resistance and could serve as antigonococcal therapeutics (17).

One key question prompted by this work is how neutrophils affect the dynamics of Gc sialylation. Neutrophils have been proposed as a source of CMP-NANA, since lysates from polymorphonuclear phagocytes, including neutrophils, confer “unstable” serum resistance on Gc (77). How Gc obtains CMP-NANA, which is cytoplasmic, is currently unknown, since intracellular Gc reside in phagolysosomes (65). Phagosomes may contain sialic acid transporters; in addition, neutrophils may release CMP-NANA by lysis or elaboration of neutrophil extracellular traps. It is also tantalizing to speculate that neutrophil sialyltransferases, found in the secretory pathway, could intersect with Gc- and sialic acid-containing compartments to sialylate the bacteria (78). Conversely, neutrophils have surface-exposed sialidases, which desialylate their own glycans during diapedesis (79, 80) and other cells’ glycans for adhesion (81). While we did not detect any desialylation of Gc over the first 30 min of neutrophil exposure, the sialylation state of Gc over time with neutrophils remains unknown. Modification of neutrophil-like cells to synthesize azido-labeled CMP-NANA (82) would allow sialylation of Gc to be tracked over time and in different

Sialylated Gc have reduced infectivity in experimental challenge of the human male urethra (14) and in the genital tract of female mice (15, 16). However, Gc isolated from male urethral gonorrheal exudates are sialylated (8, 11, 21). Additionally, Gc isolated from cervicovaginal human secretions are less likely to be sialylated, due to sialidase activity conferred by members of the female genital microbiome (83). Given these dynamics, we anticipate that balancing sialylation state allows Gc to resist cellular and humoral immunity – including by engaging Siglecs to thwart neutrophil attack – while maintaining infectivity of target mucosal surfaces. These findings can be exploited for novel therapies for gonorrhea, since Lst expression is not phase variable. For instance, CMP-NANA analogs that do not confer serum resistance aid complement killing of multidrug-resistant Gc (17, 84). Similarly, sialic acid analogs that do not engage Siglecs could be developed to render Gc more sensitive to neutrophils. These approaches, in conjunction with antibiotics and vaccines, could enhance neutrophil and soluble host defenses to combat drug-resistant gonorrhea.

## MATERIALS AND METHODS

### Bacterial strains and growth conditions

Gc in this study are in the FA1090 background, constitutively expressing the pilin variant 1-81-S2 and with in-frame deletions of all *opa* genes (Opaless) (48). Opaless Gc with non-phase variable, constitutively expressed *opaD* has been described previously (OpaD) (48); Opa60 and OpaI were constructed similarly (49, 66).

Gc were grown overnight on gonococcal medium base (Difco) with Kellogg’s supplements (85) (GCB) at 37°C with 5% CO_2_. Growth of Gc in rich liquid medium containing Kellogg’s supplements and NaHCO_3_ to enrich for piliated, live, mid-logarithmic phase bacteria has been previously described (86). For sialylation, 50µg/mL of cytidine-5’-monophosphate-N-acetylneuraminic acid (CMP-NANA) (Nacalai) reconstituted in PBS was added for the final 2.5 h growth (87). Gc were washed with PBS + 5mM MgSO_4_ prior to infection. CMP-Az-NANA (R&D Systems) was added in lieu of or in competition with unmodified CMP-NANA (54).

Isogenic OpaDΔ*lst* was generated by spot transforming OpaD with genomic DNA from MS11 GP300 *lst::*Kan (from A. Jerse, USUHS) (15). Transformants were selected on GCB with 50µg/mL kanamycin and confirmed by PCR, and inability to sialylate OpaDΔ*lst* was verified by 6B4 exclusion (see below). OpaD expression was confirmed by Western blotting bacterial lysates with 4B12 mAb against Opa (88, 89).

### Gc sialylation and detecting sialylated Gc

Western blot: SDS-PAGE and immunoblotting for 6B4 reactivity was conducted as previously described (17). In brief, lysates from Gc with or without CMP-NANA pre-incubation, were resolved using a 4-20% gradient SDS-polyacrylamide gel (Bio-Rad) and transferred in CAPS methanol buffer to polyvinylidene difluoride (PVDF). Membranes were blocked in PBS + 0.5% Tween-20 + 5% dry skim milk and probed with 6B4 tissue culture supernatant (from S. Ram, UMass) followed by anti-mouse IgM-horseradish peroxidase (HRP) antibody (Jackson ImmunoResearch). Blots were developed with SuperSignal chemiluminescent substrate (ThermoFisher). For imaging flow cytometry, Gc (10^8^ CFU/mL), incubated with or without 50µg/mL CMP-NANA, were labeled with 5 µM Tag-IT Violet (TIV) (BioLegend) then incubated with 6B4, followed by AlexaFluor647-coupled goat anti-mouse IgM (Jackson ImmunoResearch) before fixation and processing. Samples were examined using ImageStream X Mark II imaging flow cytometer and analyzed using INSPIRE and IDEAS v. 6.2 software packages (Amnis Luminex Corporation). Focused, singlet TIV+ Gc were gated as in (66). AF647+ gate was set in reference to TIV+ Gc without 6B4. Data are reported as fluorescence index (geometric mean fluorescence intensity x percent AF647+).

Click chemistry: Gc (10^8^ CFU/mL) with or without CMP-Az-NANA were stained with TIV before fixation to analyze Gc alone, or used to infect primary human neutrophils at a multiplicity of infection (MOI) of 1 (see below). After fixation with 2% paraformaldehyde (PFA), neutrophils were incubated with H.5 mouse anti-PorB (from M. Hobbs, UNC) to detect extracellular Gc, then permeabilized with 0.2% saponin. Copper-catalyzed click chemistry was carried out for 1 hr at room temperature with continuous shaking, using reaction reagents: 2.5mM THPTA (tris-hydroxypropyltriazolylmethylamine) ligand (Lumiprobe), 0.06mM FAM alkyne 5-isomer (5-Carboxyfluorescein) (Lumiprobe), 2.4 mM L-ascorbic acid (VWR), and 2mM copper (II) sulfate pentahydrate all dissolved in UltraPure distilled water (Invitrogen) (90). Samples were thoroughly washed before analysis by imaging flow cytometry.

### Neutrophil isolation

Human subjects research was conducted in accordance with a protocol approved by the University of Virginia Institutional Review Board for Health Sciences Research (#13909). Informed, formal written consent was obtained from each participant. Neutrophils were isolated from venous blood collected from healthy human subjects via dextran sedimentation followed by a Ficoll gradient as described (86). Neutrophils were resuspended in Dulbecco’s phosphate buffered saline (DPBS; without calcium and magnesium; Thermo Scientific) with 0.1% dextrose and used within 1 h after isolation. Samples were > 95% neutrophils and > 99% viable.

### Neutrophil ROS production

Neutrophils were resuspended in Morse’s Defined Medium (MDM) (91) containing 20µM luminol. Gc was added at an MOI of 100 and the generation of reactive oxygen species was measured every 3 min for 1 h via chemiluminescence (86). Negative controls included Opaless-infected and uninfected neutrophils (48). Line graphs are one representative of three or more biological replicates. Bar graphs present all biological replicates’ area under the curve (AUC) values.

### Neutrophil granule exocytosis

Surface expression of neutrophil granule proteins was measured as in (42), using phycoerythrin (PE)-CD63 for primary granules and allophycocyanin (APC)-CD66b for secondary granules (or isotype controls PE-IgG1 and APC-IgM) (BioLegend). Data were acquired using a Cytek Northern Lights spectral flow cytometer and analyzed using FCS Express (De Novo Software). The median fluorescence of each sample was normalized to unstimulated neutrophils as negative control.

### Gc survival in the presence of neutrophils

Infection of adherent, IL-8 treated human neutrophils in RPMI 1640 medium with 10% heat-inactivated fetal bovine serum (HI-FBS) was conducted as in (86), except Gc was pre-incubated with or without CMP-NANA. After centrifugation to synchronize infection and incubation at 37°C, 5% CO_2_, neutrophils were lysed with 1% saponin, serially diluted, and plated for overnight growth on GCB. Results are reported as percentage of CFUs enumerated from lysates at the indicated timepoint, divided by CFUs enumerated at 0 min.

### Imaging flow cytometric analysis of Gc association with and internalization by neutrophils

Infection of primary human neutrophils with TIV+ Gc and evaluation by imaging flow cytometry was performed as described (42, 68, 92). Because sialylation reduced bacterial binding of the goat anti-Gc antibody (Biosource) to almost undetectable levels (data not shown), extracellular Gc were recognized instead with H.5 anti-PorB, followed by AF647-coupled goat anti-mouse antibody.

### NCEACAM-1 and NCEACAM-3 binding of Opa expressing Gc

GST-tagged N-terminal domains of human CEACAMs 1 and 3 were purified and utilized as in (66). In brief, nonsialylated or sialylated Gc were incubated with GST-N-CEACAM1 or -N-CEACAM3 for 30 min at 37°C with end-over-end rotation. Gc were washed and incubated with mouse anti-GST antibody p1A12 (BioLegend) followed by an AF488-conjugated goat anti-mouse antibody (Jackson ImmunoResearch). The percentage of AF488+ (N-CEACAM-bound) Gc was measured using imaging flow cytometry.

### Gc association with CHO-hCEACAM-1 and -3 cells

hCEACAM1-, hCEACAM3-, and Control-CHOs were generated using heparan-sulfate proteoglycan deficient Chinese hamster ovary (CHO) cell strain PgsD-677 (ATCC). Cells were grown in Ham’s F-12K medium (ATCC) with HI-FBS and 1x Antibiotic-Antimycotic (Gibco) at 37°C, 5% CO_2_. Plasmids were prepared from VectorBuilder *Escherichia coli* stocks using Plasmid Miniprep (Qiagen) and ampicillin selection: human CEACAM1 (VB900000-5687rxc), human CEACAM3 (VB900000-7471zkb), or an empty control (VB010000-9288rhy) vector under a CAG promoter. CHOs were transfected using Lipofectamine 3000 transfection reagent (Invitrogen) according to manufacturer instructions. After 48 h, transfectants were selected using 10µg/mL puromycin (Gibco). A non-transfected control was used to confirm selection. Transfectants were maintained under puromycin selection throughout culturing and passaged at 70-90% confluency; medium was replaced with antibiotic-free medium 2 h prior to infection. Surface expression of CEACAM on pooled CEACAM1- and 3-CHO transfectants, and absence on Control-CHOs, were confirmed by immunofluorescence microscopy using anti-Pan-CEACAM antibody (Abcam, clone D14HD11).

Cells were seeded in a 6-well plate and grown for 48 hr to confluency (∼10^6^ cells/well). TIV+ Gc were added at an MOI of 1 to cells on ice, centrifuged to synchronize infection, washed, and incubated for 30 min. Cells were lifted with Versene solution (Gibco) at 37°C, 5% CO_2_ for 10 min. Cells were then fixed with 2% PFA, collected with a rubber cell scraper, and filtered through 50µm nylon mesh. Cells were blocked in PBS + 10% normal goat serum and incubated with anti-Pan-CEACAM antibody followed by goat anti-mouse AF555-conjugated antibody (Invitrogen). Cells were analyzed using imaging flow cytometry and IDEAS software. Focused cells were gated by gradient root mean square for image sharpness ≥50. Single cells were gated by high aspect ratio and area ≥200 and ≤1000. Single stained controls were used to set gates for AF555+ and TIV+ populations.

### Siglec staining and blockade

Adherent, IL-8 primed neutrophils were fixed and lifted with a cell scraper then cells were washed in PBS and blocked in 10% normal goat serum. Next, neutrophils were incubated with anti-Siglec-9-APC (Clone 191240) and/or anti-Siglec-5/14-AF488 (Clone 194128) (R&D Systems) for 30 min. Samples were analyzed using imaging flow cytometry as above. To block Siglecs, isolated neutrophils were incubated with the unconjugated forms of anti-Siglec-9 and/or anti-Siglec-5/14 for 30 min at 37°C + 5% CO_2_ before infection.

### Statistics

Results are depicted as the means ± standard errors for ≥ 3 independent experiments. Statistics were calculated using GraphPad Prism; a *p* value of ≤ 0.05 was considered significant. Analysis of variance (ANOVA) or mixed model analysis was used for multiple comparisons for parametric data.

## Acknowledgements

This work was supported by NIH R01AI097312, NIH U19AI144180, and NIH U01AI162457 (AKC). AJC was supported in part by NIH F31AI157528 and NIH 3R01AI097312-07S1. AJC and MWB were supported in part by NIH T32AI007046. MMP and NJF were supported by NIH R35GM124893. GAJ and CMJ were supported by Merit Review Award BX000727 and Research Career Scientist Award from the Research Service of the US Department of Veterans Affairs. The funders had no role in study design, data collection and interpretation, or the decision to submit the work for publication. We thank Louise Ball and Samuel Clark for preliminary experiments that informed this project, and Asya Smirnov and Linda Columbus for experimental advice. We thank Mike Solga of the UVA Flow Cytometry Core Facility (RRid:SCR_017829) and Hazel Ozuna (University of Louisville) for advice, Ann Jerse and Marcia Hobbs for strains and antibodies, and Sanjay Ram for reagents and discussions. We thank the human subjects who make this research possible.

## Author contributions

Conceptualization: AJC, AKC. Formal analysis: AJC, KST, NJF. Funding acquisition: AKC, AJC, GAJ, MMP. Project Administration: AKC. Investigation: AJC, KST, MWB, CMJ, NJF. Methodology: AJC, MWB, NJF, CMJ, MMP. Supervision: GAJ, MMP, AKC. Visualization: AJC. Writing – original draft: AJC, AKC. Writing – review & editing: AJC, KST, MWB, NJF, CMJ, MMP, GAJ, AKC.

